# Elucidating another level of epigenetic regulation in osteoarthritis by identifying functional *cis*-acting long non-coding RNAs and their targets in articular cartilage

**DOI:** 10.1101/2020.03.23.003020

**Authors:** Marcella van Hoolwerff, Paula I. Metselaar, Margo Tuerlings, H. Eka D. Suchiman, Nico Lakenberg, Yolande F.M. Ramos, Davy Cats, Rob G.H.H. Nelissen, Demiën Broekhuis, Hailiang Mei, Rodrigo Coutinho de Almeida, Ingrid Meulenbelt

## Abstract

**Objective:** To identify robustly differentially expressed long non-coding RNAs (lncRNAs) with osteoarthritis (OA) pathophysiology in cartilage. Moreover to explore potential target mRNAs by establishing co-expression networks, followed by functional validation.

**Methods:** RNA sequencing was performed on macroscopically lesioned and preserved OA cartilage of patients who underwent a joint replacement surgery due to OA (N=98). Differential expression (DE) analysis was performed on lncRNAs that were annotated in GENCODE and Ensembl. To identify potential interactions, correlations were calculated between the identified DE lncRNAs and previously reported DE protein-coding genes in the same samples. Modulation of chondrocyte lncRNA expression was achieved using LNA GapmeRs.

**Results:** By applying our in-house pipeline we identified 5,053 lncRNAs to be robustly expressed, of which 191 were FDR significant differentially expressed between lesioned and preserved OA cartilage. Upon integrating mRNA sequencing data, we showed that intergenic and antisense DE lncRNAs show high, positive correlations with their flanking, respectively, sense genes. To functionally validate this observation we selected *P3H2-AS1*, which was downregulated in primary chondrocytes, resulting in downregulation of *P3H2* gene expression levels. As such, we can confirm that *P3H2-AS1* regulates its sense gene *P3H2*.

**Conclusion:** By applying an improved detection strategy, robustly differentially expressed lncRNAs in OA cartilage were detected. Integration of these lncRNAs with differential mRNA expression levels in the same samples showed insight into their regulatory networks. Our data signifies that intergenic, as well as antisense lncRNAs play an important role in regulating the pathophysiology of OA.

## INTRODUCTION

Osteoarthritis (OA) is an age-related, heterogenous, degenerative disease of the articular joints, characterized by, amongst others, cartilage degeneration and remodeling of subchondral bone, resulting in stiff and painful joints and decreased mobility [1]. Despite the fact that OA is the most prevalent of joint diseases worldwide, no effective treatment is available at the moment [2]. It has been shown that pathophysiology of OA in cartilage is marked by altered gene expression regulation in chondrocytes [3, 4]. This alteration of gene expression regulation could be triggered by adaptation processes occurring due to aging, genetic predisposition or environmental stimuli and is in part caused by aberrant epigenetic mechanisms. These mechanisms include DNA methylation, histone modifications and expression of microRNAs (< 22 nucleotides) [4–6]. More recently, long non-coding RNAs (lncRNAs, > 200 nucleotides) have been shown to play an important role in the homeostasis of the extracellular matrix of cartilage [5, 7–10]. LncRNAs are defined as RNA transcripts with little or no protein-coding potential and are known to regulate transcription and translation by numerous mechanisms, such as chromatin remodeling, mRNA stabilization, microRNA modulation, and recruitment of scaffolding proteins. One type of classification of lncRNAs is based on the genomic location with respect to protein-coding genes, so-called biotypes, including antisense RNAs, sense RNAs, pseudogenes, and long intergenic non-coding RNAs (lincRNAs). Another type of classification is based on the location at which the lncRNA functions relative to its transcription site, which can be in *trans* or *cis* [11–13]. *Cis*-acting lncRNAs comprise a considerable portion of known lncRNAs and can be positioned at various distances and orientations relative to their target genes, such as lincRNAs around transcription factor start sites, as well as sense and antisense lncRNAs that overlap with their sense gene [13, 14]. Potentially, lncRNAs could be candidate targets in OA treatment, since their expression can be highly tissue specific [9].

RNA sequencing (RNA-seq) has made improvements possible in detecting lncRNAs, nevertheless, mapping and annotating lncRNAs remains challenging. These challenges arise from the fact that they are usually very lowly expressed and their sequence-function relationship is still poorly understood. Moreover, recent ribosome profiling and bioinformatics studies have suggested that a large proportion of transcripts is with unknown coding potential [15]. Recent studies with respect to OA have focused on intergenic lncRNAs, even though the fraction genic and intergenic lncRNAs can be equal in size depending on the tissue investigated [15, 16]. To determine the complete lncRNA transcriptome, we applied an in-house pipeline to robustly capture lncRNAs in a previously assessed RNA-seq dataset of lesioned and preserved OA cartilage samples [4]. Subsequently, lncRNAs associated with OA pathophysiology were identified and potential interactions with OA-specific mRNAs were investigated.

## MATERIALS AND METHODS

### Sample collection

Macroscopically lesioned and preserved articular cartilage samples were obtained from the Research osteoArthritis and Articular Cartilage (RAAK) study as described in Ramos *et al*. [3]. In this study, a total of 98 samples was used (65 knees, 33 hips) (**online supplementary table 1**). Ethical approval was obtained from the medical ethics committee of the LUMC (P08.239/P19.013) and informed consent was obtained from all participants.

### RNA sequencing

Total RNA from articular cartilage was isolated using Qiagen RNeasy Mini Kit (Qiagen, GmbH, Hilden, Germany). Paired-end 2×100 bp RNA sequencing (Illumina TruSeq RNA Library Prep Kit, Illumina HiSeq2000 and Illumina HiSeq4000) was performed. Strand specific RNA-seq libraries were generated which yielded a mean of 20 million reads per sample. Quality control was performed as described in [4], subsequently reads were aligned to the GRCh38 reference genome with the RNA-seq aligner STAR (v2.6.0) [17]. Hereafter, aligned reads were processed into individual transcripts using StringTie (v1.3.4) [18]. LncRNAs were identified by mapping the transcripts to GENCODE (v29) [11] and Ensembl (v94) [19].

In order to filter out transcripts with unknown coding potential, we integrated two sources of evidence: (1) predictions from the alignment-free Coding Potential Assessment Tool (CPAT, v1.2.2) and (2) predictions from the LncFinder R package (v1.1.3). CPAT is a machine learning based method which analyzes the sequence features of transcript open read frames (ORFs) using a logistic regression model built from ORF size, Fickett TESTCODE statistic and hexamer usage bias. LncFinder predicts lncRNAs using heterologous features and machine learn model [20]. Transcripts with coding potential predicted by both tools were removed from the dataset.

### Differential expression analysis and replication

Differential expression (DE) analysis was performed on 32 paired samples (25 knees and 7 hips) (**online supplementary table 1A**) using the DESeq2 R package (v1.24) [21]. We applied a general linear model assuming a negative binomial distribution, followed by a paired Wald-test between lesioned and preserved OA cartilage samples, where the preserved samples were set as the reference. Benjamin-Hochberg multiple testing corrected P-values were considered significant at < 0.05 and are reported as false discovery rate (FDR). Furthermore, to validate the results, five significant DE lncRNAs were selected and measured by RT-qPCR in 10 paired cartilage samples overlapping with the RNA-seq samples (**online supplementary table 1B**), replication was performed in an independent cohort of 10 paired cartilage samples (**online supplementary table 1C**). Total RNA was isolated using RNeasy Mini Kit (Qiagen), followed by cDNA synthesis using 100 ng RNA with the First Strand cDNA synthesis kit (Roche Applied Science) according to manufacturer’s protocol. Expression levels were determined using FastStart SYBR Green Master reaction mix (Roche Applied Science) for *AC025370.1, AC090877.2, MEG3, P3H2-AS1, TBILA*, and *GAPDH*. Primer sequences are shown in **online supplementary table 2**. Relative gene expression levels were calculated with the 2^−ΔΔCt^ method, using *GAPDH* as internal control. A paired t-test was performed on the -ΔCt values, and P < 0.05 was considered significant.

### LncRNA-mRNA interactions

LncRNA expression data was normalized and variance stabilizing transformed (vst) using the DESeq2 R package (v1.24) [21] and batch effect was removed using the limma R package (v3.40.6) [22]. Our previously published mRNA data [4] was equally normalized and transformed, and batch effect was equally removed. Subsequently, Spearman’s correlations were calculated between the FDR significant DE lncRNAs and the DE protein-coding genes previously published [4] using the Hmisc R package (v4.2.0) in OA cartilage samples (**online supplementary table 1D**). Correlations with P < 0.05 were considered significant.

### *In vitro* downregulation lncRNA using locked nucleic acid GapmeRs

Primary chondrocytes were isolated from three independent donors and passaged twice or thrice, as described in [23]. Chondrocytes were transfected in duplo with antisense locked nucleic acid (LNA) GapmeR (Qiagen) targeting *P3H2-AS1* (TGAGCAACTAGGTGTA) or GapmeR negative control (AACACGTCTATACGC) at 10 nM final concentration using Lipofectamine RNAiMax Transfection Reagent (Invitrogen) according to manufacturer’s protocol. Cells were lysed 30 hours post transfection with TRIzol Reagent (Thermo Fisher Scientific) for RNA isolation, which was done using RNeasy Mini Kit (Qiagen). Synthesis of cDNA was performed with 150 ng of total RNA using the First Strand cDNA Synthesis kit (Roche Applied Science) according to the manufacturer’s protocol. Expression levels were determined using FastStart SYBR Green Master reaction mix (Roche Applied Science) for *P3H2-AS1, P3H2*, and *GAPDH*. Primer sequences are shown in **online supplementary table 2**. Relative gene expression levels were calculated with the 2^−ΔΔCt^ method, using *GAPDH* as internal control. A paired t-test was performed on the -ΔCt values, P < 0.05 was considered significant.

## RESULTS

### Characterization of lncRNAs in OA cartilage

To characterize lncRNAs in OA cartilage, we used our previously assessed RNA-seq data of 32 paired samples (25 knees, 7 hips, **online supplementary table 1A**) of lesioned and preserved OA cartilage [4]. Our in-house pipeline was applied to capture lncRNAs from two different databases (GENCODE and Ensembl). As shown in **figure 1A**, initially 30,354 lncRNAs were detected in our dataset. To filter out possible transcripts of unknown coding potential, we integrated results from two machine learning approaches (CPAT and Lncfinder, see methods). After removing these transcripts the dataset remained with 29,219 lncRNAs, which were considered for further analyses. To robustly detect cartilage specific lncRNAs, a cut-off of minimal two counts on average per lncRNA was applied, resulting in a total of 5,053 lncRNAs expressed in cartilage (**figure 1A**). Classification of these lncRNAs based on biotype, showed that 1,989 were antisense RNAs (39.4%), 249 sense RNAs (4.9%), 1,532 pseudogenes (30.3%), and 900 lincRNAs (17.8%) (**figure 2**).

**Figure 1.**
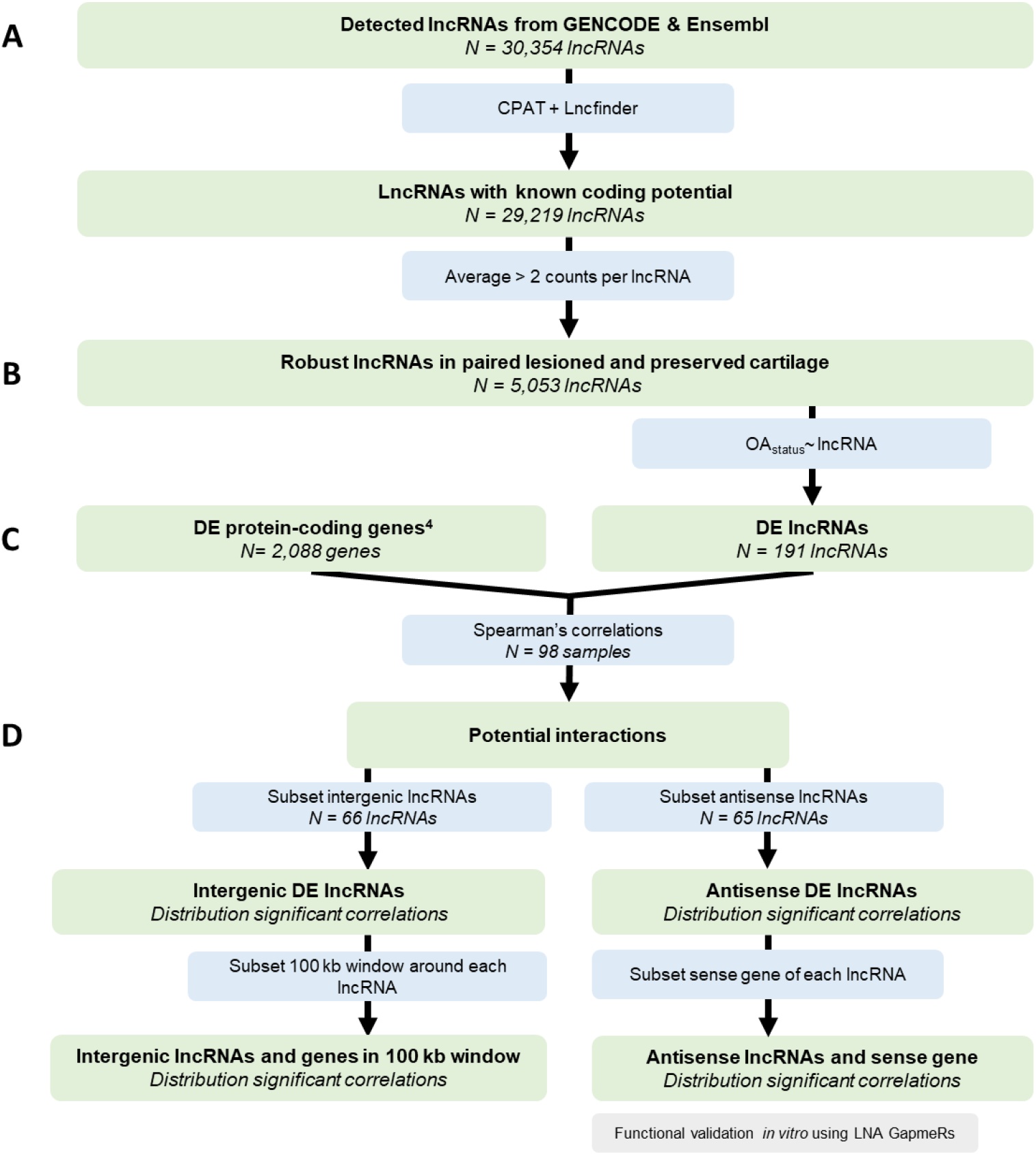
Overview of applied strategy. Number of genes or lncRNAs represent FDR significant genes or lncRNAs. LncRNA = long non-coding RNA, DE = differentially expressed, OA = osteoarthritis

### Differential expression lncRNAs between lesioned and preserved OA cartilage

To identify lncRNAs associated with the OA process, differential expression analysis was performed between paired lesioned and preserved OA cartilage samples, resulting in 191 significant differentially expressed (DE) lncRNAs (FDR<0.05, **figure 1B**). Of these, 65 were antisense RNAs (34.0%), 10 were sense RNAs (5.2%), 33 were pseudogenes (17.3%), and 66 were lincRNAs (34.6%) (**online supplementary table 3**). When comparing the biotypes of the total expressed lncRNAs to the biotypes of the DE lncRNAs (**figure 2**), we observed an increase of lincRNAs and a decrease of pseudogenes, suggesting that lincRNAs may be more important in the OA process. The most significant DE lncRNA was lincRNA *AL139220.2* (fold change(FC)=2.2, FDR=2.0×10^10^). As depicted in **figure 3**, 114 lncRNAs were downregulated and 77 were upregulated, with FCs ranging from 0.3 (*AC100782.1*, FDR=6.5×10^−4^) to 4.5 (*LINC01411*, FDR=2.6×10^−6^). The 191 identified lncRNAs in this study included several previously found to be associated to OA, such as *MEG3* (FC=0.6, FDR=8.8×10^−3^), *PART1* (FC=1.8, FDR=1.7×10^−4^), and *LINC01614* (FC=2.6, FDR=9.5×10^−3^) [16, 24], as well as novel associated lncRNAs, including *P3H2-AS1* (FC=2.7, FDR=4.1×10^−4^) and *AC090877.2* (FC=0.3, FDR=6.2×10^−5^). Notably, previously identified lncRNAs such as *MALAT1* (FC=1.3, FDR=0.4) [25], *TUG1* (FC=1.1, FDR=0.7) [26], *HOTAIR* (FC=0.8, FDR=0.5), and *GAS5* (FC=1.1, FDR=0.8) [27] were not found to be FDR significant differentially expressed in our study.

**Figure 2.**
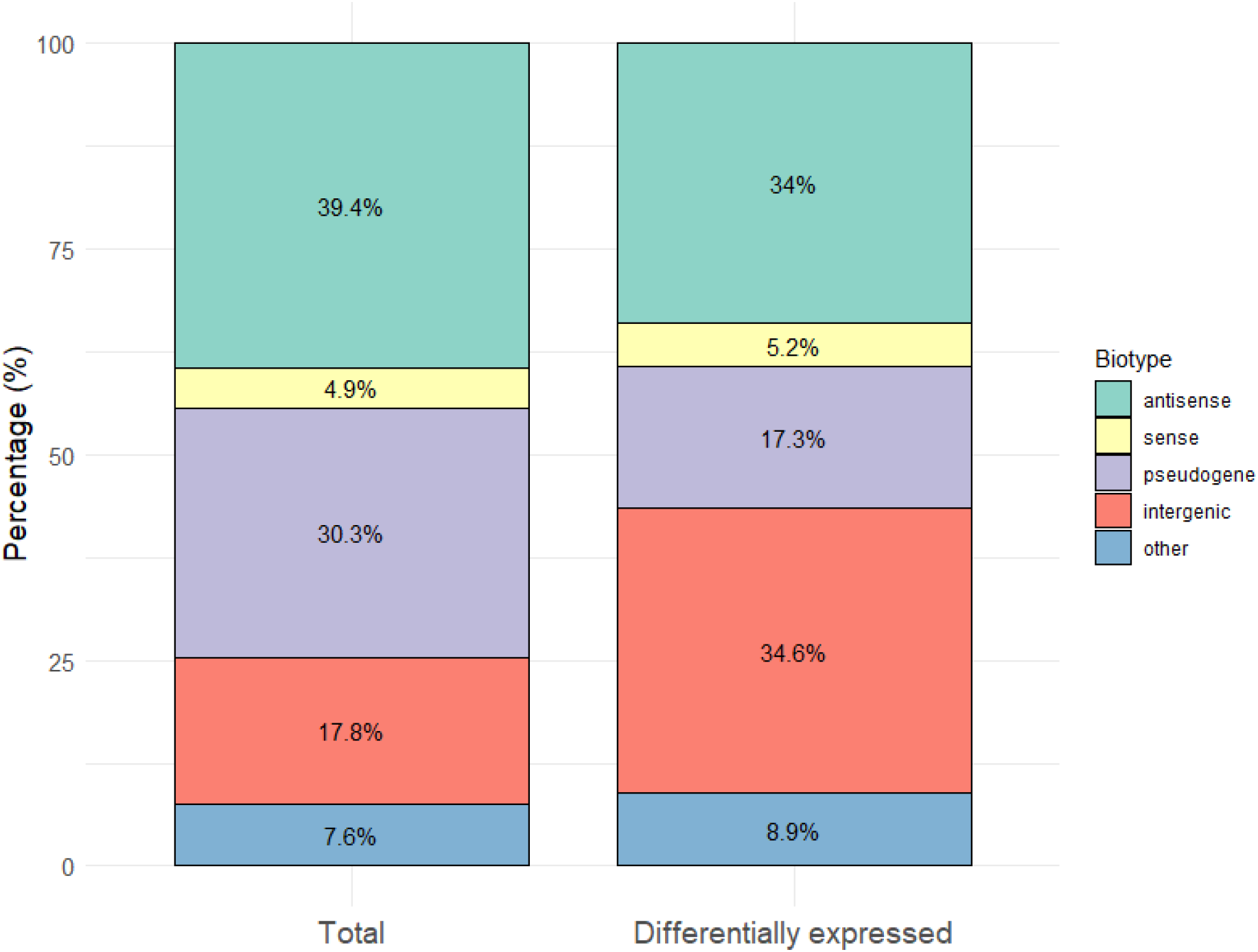
Distribution of biotypes of total lncRNAs expressed in cartilage compared to FDR significant differentially expressed lncRNAs between lesioned and preserved OA cartilage. FDR = false discovery rate, lncRNA = long non-coding RNA, OA = osteoarthritis.

**Figure 3.**
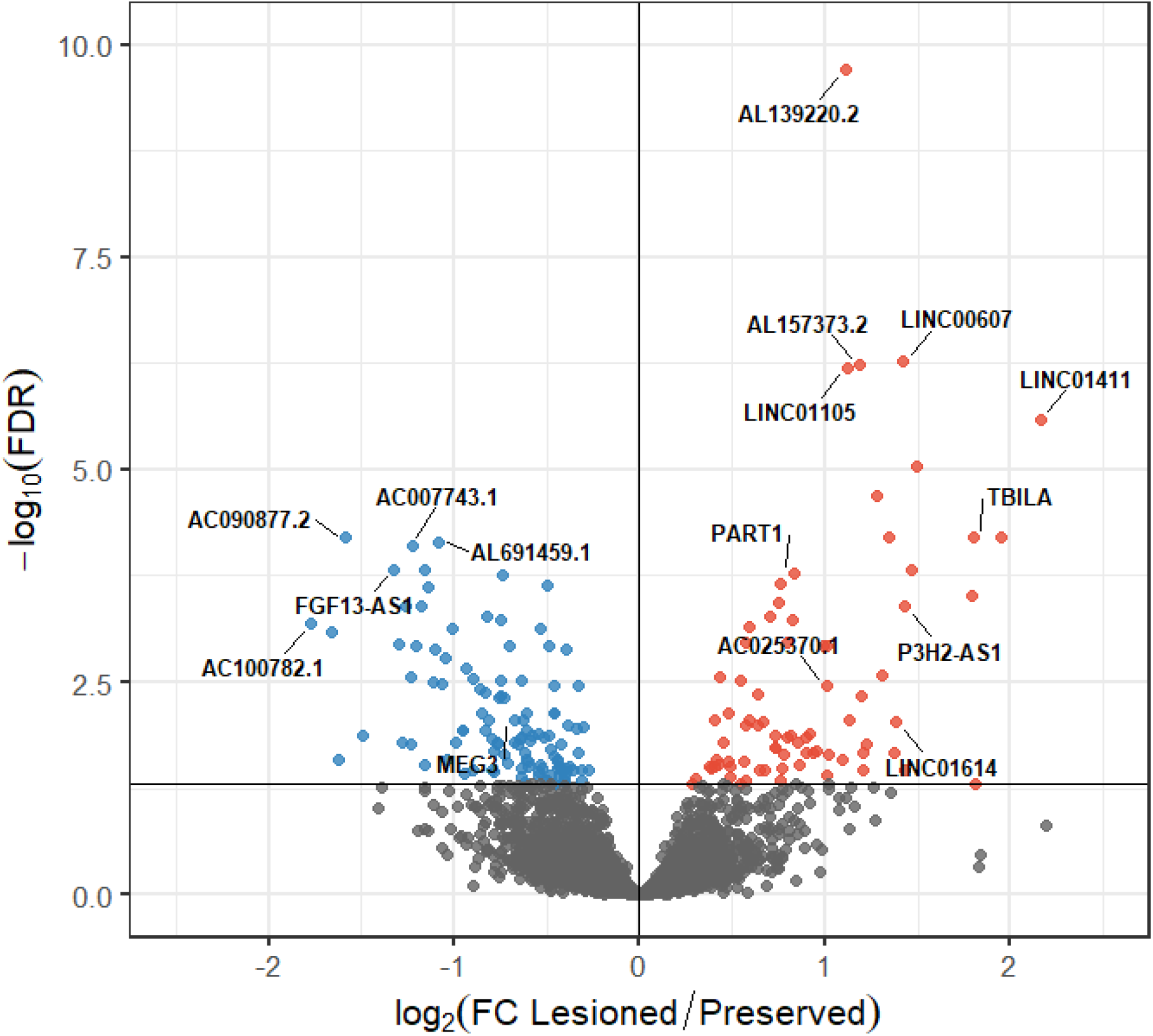
Differential expression analysis of lncRNAs between lesioned and preserved OA cartilage. Volcano plot with the differentially expressed lncRNAs, downregulated lncRNAs are depicted as blue circles and upregulated lncRNAs as red circles. Labelled are top differentially expressed lncRNAs, as well as known and novel OA associated lncRNAs. FC = fold change, FDR = false discovery rate, lncRNA = long non-coding RNA, OA = osteoarthritis.

To validate the DE results, we selected five lncRNAs (*AC025370.1, AC090877.2, MEG3, P3H2-AS1*, and *TBILA)* based on highest absolute FC and genomic location, by means of RT-qPCR in a cohort consisting of 10 paired samples (**online supplementary table 1B**) overlapping with the RNA-seq samples. All five lncRNAs were detected by RT-qPCR with equal direction of effect as those found in the RNA-seq analysis (**online supplementary table 4**). Furthermore, replication was performed in an independent cohort of 10 paired cartilage samples (**online supplementary table 1C**), which also showed comparable effect sizes and directions (**online supplementary table 4**).

### Potential interactions between lncRNAs and mRNAs relevant in OA pathophysiology

We next aimed to uncover whether mRNAs that associate with the OA process are regulated by the differentially expressed lncRNAs. Based on the assumption that interactions of lncRNAs and mRNAs likely show co-expression [28] among lesioned and preserved OA cartilage samples, correlations were calculated between our previously reported DE protein-coding genes [4] and DE lncRNAs (**online supplementary table 1D**), as shown in **figure 1C**. This resulted in 343 significant correlations (r > 0.8, **online supplementary table 5**), comprising 47 unique lncRNAs, of which 17 were antisense (36%) and 14 were intergenic (30%) (**online supplementary table 6**). These fractions are comparable to the fractions of all DE lncRNAs (**figure 2**), supporting that lncRNAs regulate mRNAs, independent of biotype. Notably, the most significant DE lncRNA, *AL139220.2* (FC=2.2, FDR=2.0×10^10^), showed one of the highest correlations with *COL6A3* (r=0.8, P=2.2×10^−16^), encoding a collagen VI chain. To visualize these interactions, an OA-specific lncRNA-mRNA co-expression network was generated. As shown in **figure 4**, three relative large clusters of interacting lncRNAs and mRNAs were observed. One cluster is characterized by being highly interlinked with a cluster of the same genes (e.g *ITGB1BP1* correlated to the six lncRNAs *IER-AS1, AL355075.3, AC234917.1, AC091564.4, AC108449.3, AL450306.1)*, whereas the other two are characterized by lncRNAs interlinked with mostly unique genes (e.g. *LNCSRLR* with 18 genes). Next to the clusters, there are a number of singular interlinked lncRNAs, one of which being *AC090877.2* (FC=0.3, FDR=6.2×10^−5^) with *GREM1* (r=0.9, P=2.2×10^−16^), which encodes a cytokine of the BMP antagonist family (**figure 4**). Interestingly, *GREM1* is the gene located closest to *AC090877.2*, suggesting that this lncRNA is *cis-*regulating this gene.

**Figure 4.**
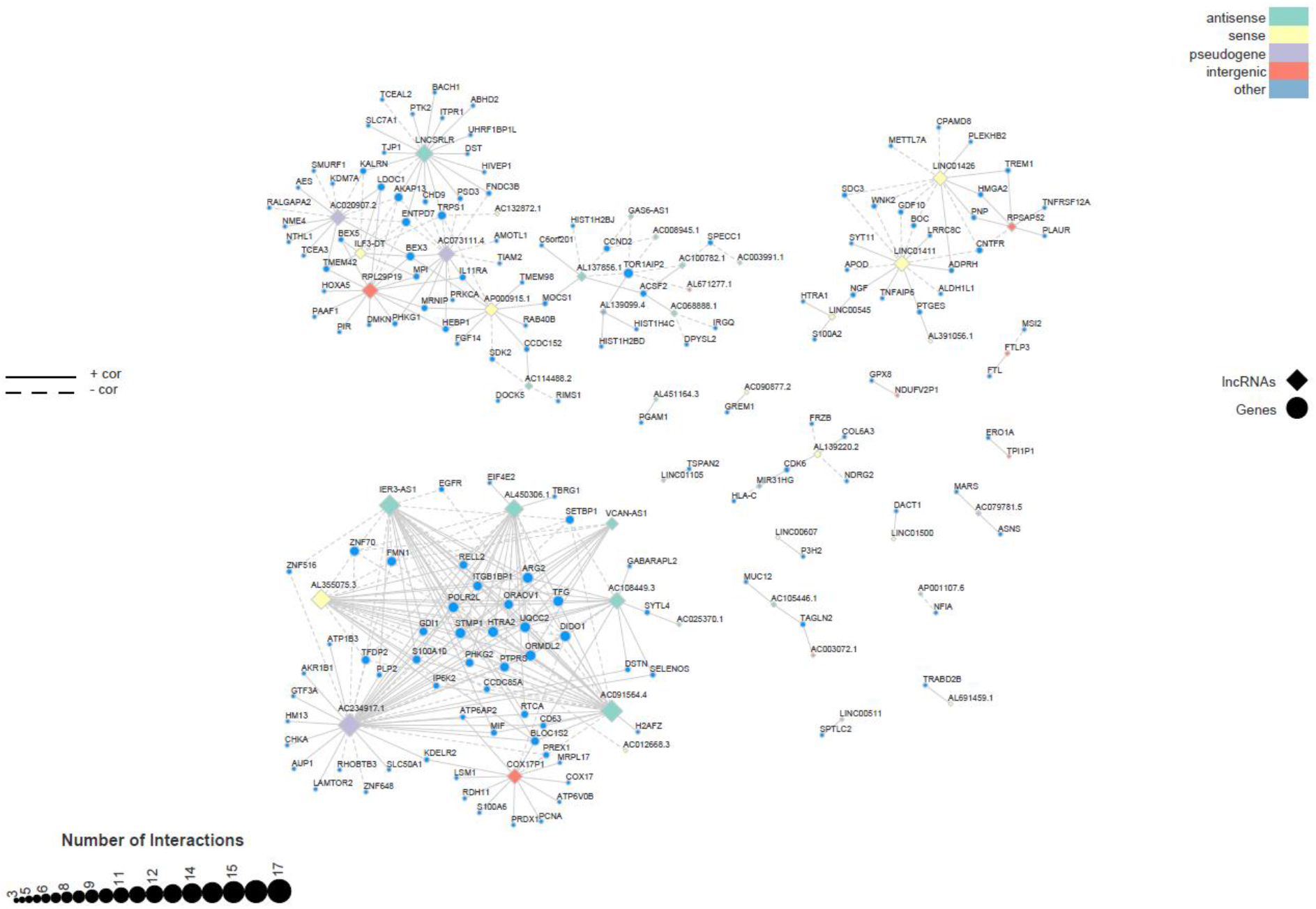
OA-specific lncRNA-mRNA co-expression network. Network of differentially expressed lncRNAs and mRNAs with correlations higher than 0.8 between preserved and lesioned OA cartilage. Diamonds are lncRNAs and the circles are mRNAs. Diamant colour characterizes the biotype of the lncRNA. The size of the node is proportional to the number of interactions. A solid line indicates a positive correlation, whereas a dashed line indicates a negative correlation. OA = osteoarthritis, lncRNA = long non-coding RNA.

The following objective was to generalize the identification of potential *cis*-regulation of DE lincRNAs (**figure 1D**). As shown in **figure 5A**, we compared the distribution of significant correlations of DE lincRNAs with all genes and DE lincRNAs with genes that lie within a 100 kb window of the transcription start site. The fraction of significant correlations > 0.5 with all DE genes was 11%, but this fraction increased to 44% when we only considered the 100 kb window. Since the fraction DE antisense lncRNAs (34%) is almost as big as the intergenic fraction (34.6%), we also aimed to identify potential *cis*-regulation of antisense lncRNAs. To this end, we compared the distribution of correlations of DE antisense lncRNAs with all protein-coding mRNAs and DE antisense lncRNAs with their sense gene (**figure 5B**). The fraction of correlations > 0.5 was 10% with all genes and 61% with only the sense genes, showing that there is an enrichment for high, positive correlations between antisense lncRNAs and their sense gene. Taken together, this data suggests that both intergenic and antisense lncRNAs are prone to regulate mRNAs in *cis* in OA cartilage.

**Figure 5.**
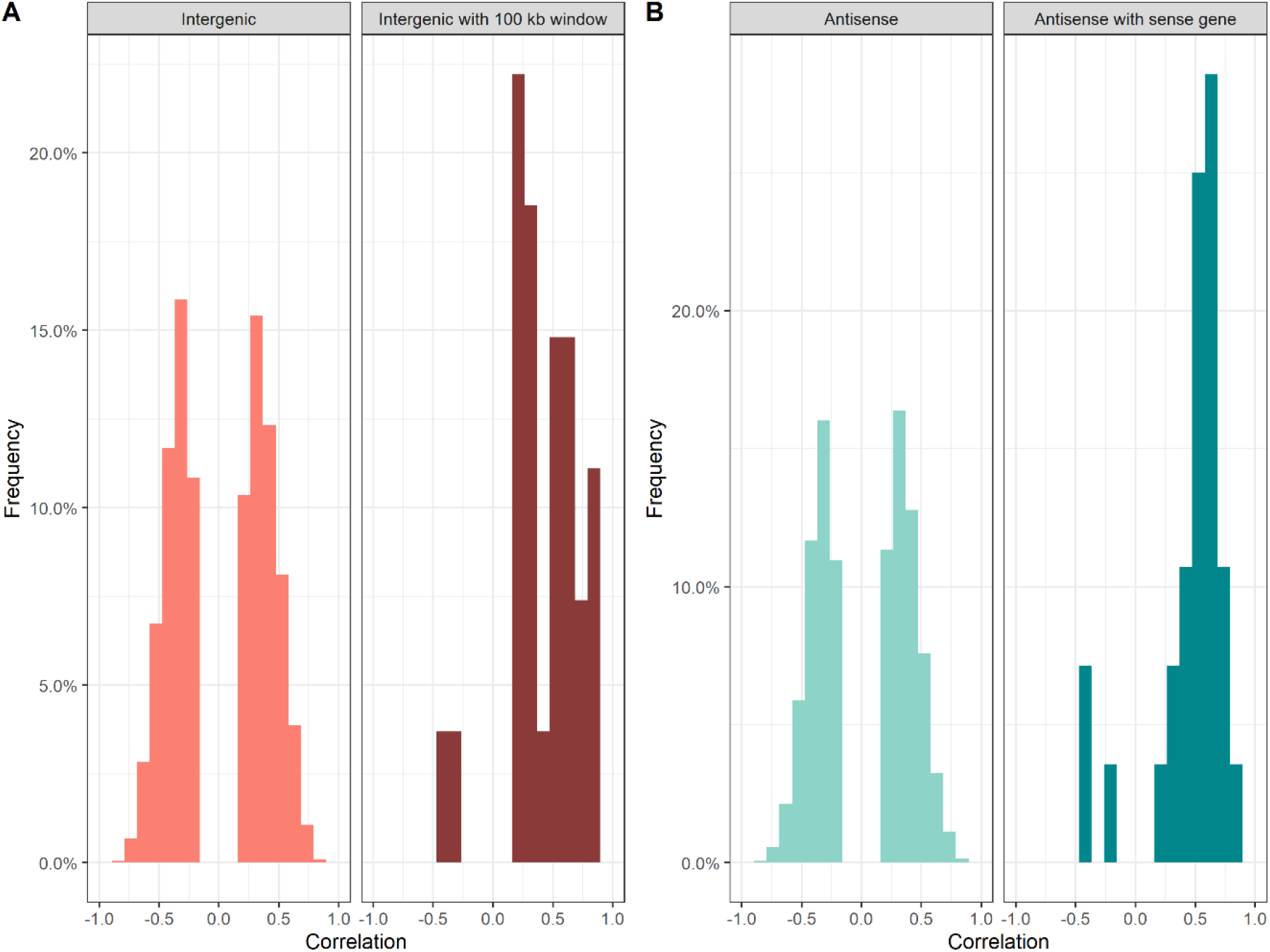
Distribution of significant correlations between intergenic DE lncRNAs (A) with previously identified DE protein-coding genes or protein-coding genes in a 100 kb window, and between antisense DE lncRNAs (B) with DE protein-coding genes or their sense genes. Correlations were calculated between lncRNA and mRNA data from the same OA cartilage samples (n=98). DE = differentially expressed, lncRNA = long non-coding RNA.

### Downregulation of lncRNA expression using locked nucleic acid GapmeRs

To validate whether the previously identified *cis*-regulation between lncRNAs and their surrounding genes is caused by a direct effect, *P3H2-AS1* was selected as a proof of concept for functional validation. *P3H2-AS1* is an antisense lncRNA, which was found to be highly upregulated in lesioned OA cartilage (FC=2.7, FDR=4.1×10^−4^) [4] and the highest correlation was with its sense gene *P3H2* (r=0.63, P=1.0×10^−13^, **online supplementary figure 1**). To this end, primary chondrocytes were transfected with a *P3H2-AS1* targeting locked nucleic acid (LNA) GapmeRs. As shown in **figure 6A**, this resulted in a significant downregulation of *P3H2-AS1* compared to a non-targeting LNA GapmeR (FC=0.28, P=0.0035). Subsequently, *P3H2* expression levels were measured, which showed that *P3H2* expression was significantly downregulated compared to cells transfected with control, non-targeting, LNA GapmeRs (FC = 0.36, P = 0.001) (**figure 6B**).

**Figure 6.**
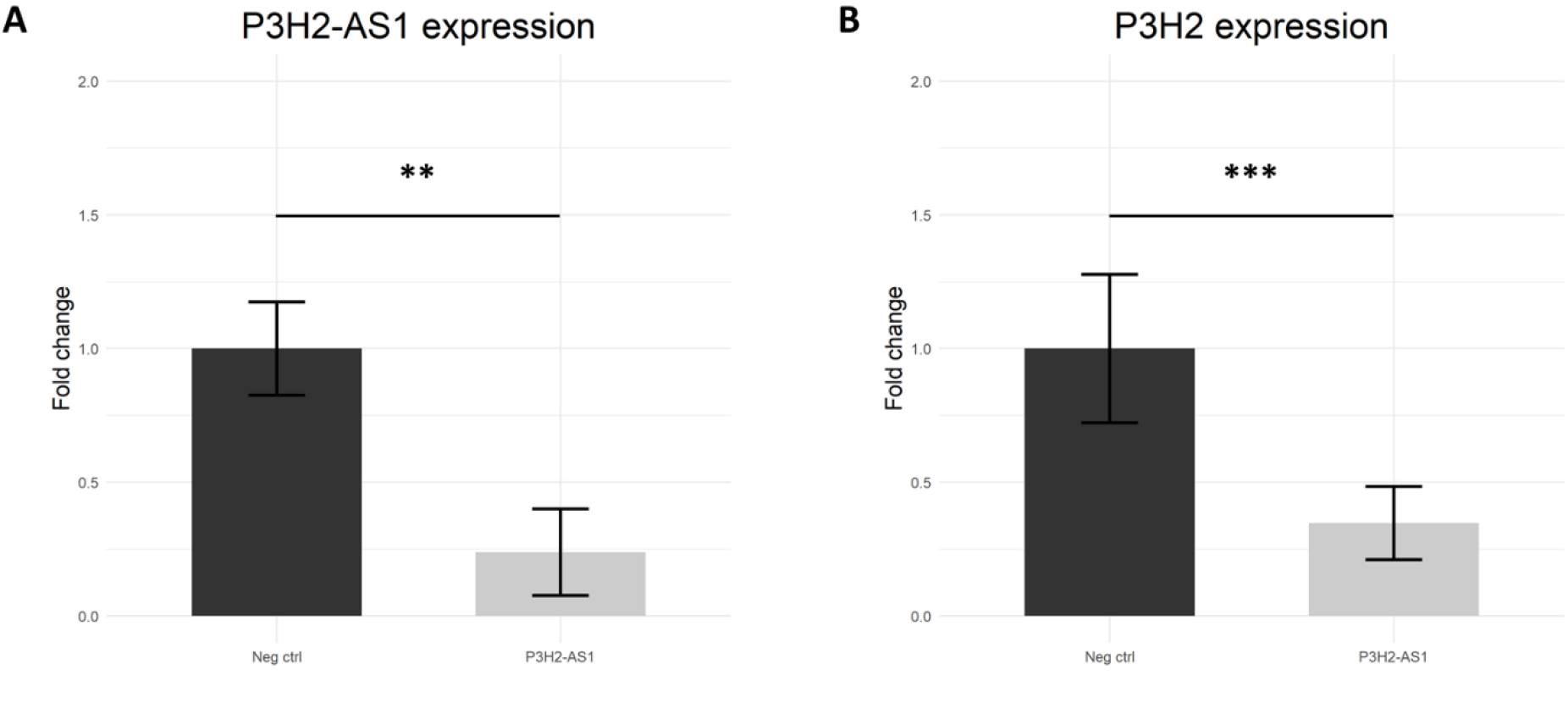
Expression of lncRNA *P3H2-AS1* (A) and gene *P3H2* (B) in primary chondrocytes transfected with *P3H2-AS1* targeting antisense LNA GapmeRs compared to non-targeting LNA GapmeRs. (A) *P3H2-AS1* expression was efficiently downregulated by the *P3H2-AS1* targeting LNA GapmeRs (B) *P3H2* expression was significantly downregulated in chondrocytes transfected with *P3H2-AS1* targeting LNA GapmeRs. Bars show mean ± SD, ** P < 0.01 *** P < 0.001 by paired t-test, n = 3 donors. lncRNA = long non-coding RNA, LNA = locked nucleic acid.

## DISCUSSION

To our knowledge, we are the first to report on robust differential expression of lncRNAs with pathophysiology of OA while integrating them with differential mRNA expression levels of the same samples using RNA sequencing. As a result, our new in-house pipeline identified 5,053 lncRNAs, robustly expressed, of which 191 were FDR significant differentially expressed lncRNAs between lesioned and preserved OA cartilage. Notably, we observed an increase in the percentage of lincRNAs, highlighting their general involvement in the OA pathophysiology process. Direction of effect of *AC025370.1* (FC=2.0, FDR=3.5×10^−3^), *AC090877.2* (FC=0.3, FDR=6.2×10^−5^), *MEG3* (FC=0.63, FDR=8.8×10^−3^), *P3H2-AS1* (FC=2.7, FDR=4.1×10^−4^), and *TBILA* (FC=3.5, FDR=1.1×10^−7^) was validated and replicated by RT-qPCR, indicating robustness of our lncRNA mapping strategy. Correlations were calculated to identify potential interactions between expression levels of DE lncRNAs and DE proteincoding genes [4] in the same OA cartilage samples. As a result, both intergenic and antisense DE lncRNAs showed an enrichment for higher, positive correlations with their flanking, respectively, sense gene compared to the total dataset. To validate this *cis*-regulation *in vitro*, *P3H2-AS1* levels were downregulated in primary chondrocytes, which resulted in downregulation of the sense gene *P3H2* expression levels, thereby confirming that *P3H2-AS1* regulates its sense gene *P3H2*.

We identified 29,219 lncRNAs to be expressed in OA cartilage, however after applying a filter with a cut-off of minimal 2 counts on average per lncRNA, the detected lncRNAs were reduced by ∼83% to 5,053. Since lncRNAs are known to be very lowly expressed, this was to be expected. However, lowly expressed lncRNAs can still be functional [12]. To allow exploratory analyses with such lowly expressed lncRNAs, deeper sequencing would be necessary, with a read-dept of e.g. 50 million reads per sample.

We showed a particular enrichment of lincRNAs in the differential expression analysis (17.8% to 34.6% **figure 2**), showing that lincRNAs indeed play an important role in the OA pathophysiology as seen in previous studies [8,16, 28]. Nonetheless, in comparison to the fraction of FDR significant DE lncRNAs reported by Pearson *et al*. [8], this fraction is still relative small. However, Pearson *et al*. performed RNA-seq on samples from isolated chondrocytes in contrast to the RNA isolated from cartilage in our study, as well as focused on profiling lncRNAs upregulated by IL-1β. The activation of chondrocyte proliferation in tissue culture will likely induce expression of RNAs involved in transcriptional regulation, as compared to the transcriptome of maturational arrested chondrocytes residing in cartilage.

Of the 191 FDR significant DE lncRNAs between lesioned and preserved OA cartilage (**figure 3**), multiple lncRNAs were previously found, including *MEG3*, *LINC01614*, and *PART1* [16, 24]. Nonetheless, there were also examples of lncRNAs previously associated to OA, which were expressed but not FDR significantly different, such as *MALAT1*, *HOTAIR*, *GAS5*, and *TUG1* [25–27]. A possible explanation could be that they were found to be DE between preserved OA and healthy cartilage, implicating they are involved in the early phase of OA pathophysiology, as opposed to genes changing during the OA process in our analysis [7]. At least 35 DE lncRNAs in our dataset were previously associated with OA [10,16, 28], however, the most significant DE lncRNA, *AL139220.2*, and the most up- and downregulated DE lncRNAs, *LINC01411* and *AC100782.1* respectively, have not been previously associated to OA, showing that a paired study design allows for the detection of many more lncRNAs involved in the OA pathophysiological process [3].

Unlike conserved microRNAs, it is difficult to predict the function of lncRNAs based solely on nucleotide sequence, due to their lack of conservation of the primary sequence [15]. To explore potential regulatory interactions between lncRNAs and mRNAs in cartilage, correlations were calculated between DE lncRNAs and DE protein-coding mRNAs (**figure 4**). At the transcriptional level lncRNAs can exert their function in *trans* or *cis* [13], which we both observed in our data. The most significant DE lncRNA, *AL139220.2*, showed one of the highest correlations with *COL6A3* (r=0.8, P=2.2×10^−16^), encoding one of the collagen VI chains as part of the complete collagen VI molecule, which is mostly present in the pericellular matrix of cartilage. *AL139220.2* is located on chromosome 1 and presently little is known about its function. Since *COL6A3* is located on chromosome 2, it seems likely that *AL139220.2* regulates *COL6A3* expression in *trans*. Markedly, *AC090877.2* showed the highest correlation with its sense gene *GREM1* (r=0.9, P=2.2×10^−16^), suggesting that this lncRNA is *cis*-regulating this gene. In previous studies, it was shown that often lincRNAs regulate flanking mRNAs in *cis* in OA, where a positive correlation was found between the expression of mRNA-flanking lincRNAs and their nearest coding mRNA [8, 28]. This observation was confirmed in our data, since the fraction of high, positive correlations (r>0.5) was considerably larger between lincRNAs and the DE genes that lie within a 100 kb window (44%) than with all DE genes (10%) (**figure 5A**). Furthermore, it is also known that antisense lncRNAs can regulate their overlapping sense genes in *cis* [14], which has not been investigated in OA previously. We found an enrichment for high, positive correlations between antisense DE lncRNAs and their sense gene (r>0.5 57%) compared to correlations between antisense DE lncRNAs and all DE genes (r>0.5 9%), suggesting that indeed antisense lncRNAs often regulate their sense gene in *cis* (**figure 5B**). Hence, to completely understand the transcriptional regulation of lncRNAs in the OA process, the total lncRNA transcriptome should be considered and not solely the lincRNAs. Of importance is the notion that these correlations are not yet proof of a (direct) downstream effect of lncRNAs on the mRNAs.

Given these observations, we selected the antisense lncRNA *P3H2-AS1* as proof of principle to establish whether it regulates its sense gene. Downregulation of *P3H2-AS1* resulted in a significant downregulation of *P3H2* expression levels, thereby confirming that *P3H2-AS1* regulates its sense gene in *cis* (**figure 6**). *P3H2* encodes an enzyme that catalyzes post-translational 3-hydroxylation of proline residues and plays a critical role in collagen chain assembly, stability, and cross-linking and was recently found to be highly upregulated in lesioned OA cartilage and therefore likely involved in the OA process [4]. Antisense lncRNAs can affect biogenesis or mobilization of target RNA on multiple levels, such as transcription, splicing, and translation [14]. To elucidate the exact mechanism of *P3H2-AS1* regulating *P3H2*, complementary functional studies employing e.g. CRISPR/Cas9, RNA fluorescence *in situ* hybridization, or crosslinked immunoprecipitation are necessary, to be able to investigate whether *P3H2-AS1* can be used as a potential preclinical target by modulating *P3H2* expression levels via *P3H2-AS1* [29].

In conclusion, our improved detection strategy resulted in characterization of lncRNAs robustly expressed in OA cartilage. Our data signifies that intergenic, as well as antisense lncRNAs play an important role in regulating the pathophysiology of OA. Moreover, we observed that besides the previously found notion that intergenic lncRNAs function in *cis*, antisense lncRNAs can also exert their function in *cis*, which we confirmed *in vitro*. Future studies regarding lncRNAs and OA should be complemented by functional validation, e.g. by modulating lncRNA expression levels using antisense LNA GapmeRs, to confirm whether a correlation equals a biological relation between lncRNA and mRNA.

## ACKNOWLEDGEMENTS

We thank all the participants of the RAAK study. The LUMC has and is supporting the RAAK study. We thank Enrike van der Linden, Robert van der Wal, Peter van Schie, Shaho Hasan, Maartje Meijer, Daisy Latijnhouwers, and Geert Spierenburg for their contribution to the collection of the joint tissue.

## Supplementary materials

**Supplementary table 1** - Sample characteristics included in the analysis in the (A) discovery, (B) validation (C) replication, and (D) correlation analyses.

**Supplementary table 2** - Primer sequences to measure mRNA and lncRNA expression levels.

**Supplementary table 3** - FDR significant differentially expressed lncRNAs in lesioned versus preserved OA cartilage samples

**Supplementary table 4** - Results of the validation and replication of DE lncRNAs between lesioned and preserved OA cartilage, paired t-test was performed on the -ΔCt values

**Supplementary table 5** - Nominal significant correlations between FDR significant differentially expressed lncRNAs and FDR significant differentially expressed protein-coding genes in the same OA cartilage samples

**Supplementary table 6** - Count and percentage of unique lncRNAs that have correlations > 0.8 with DE mRNAs from the same samples

**Supplementary figure 1.**
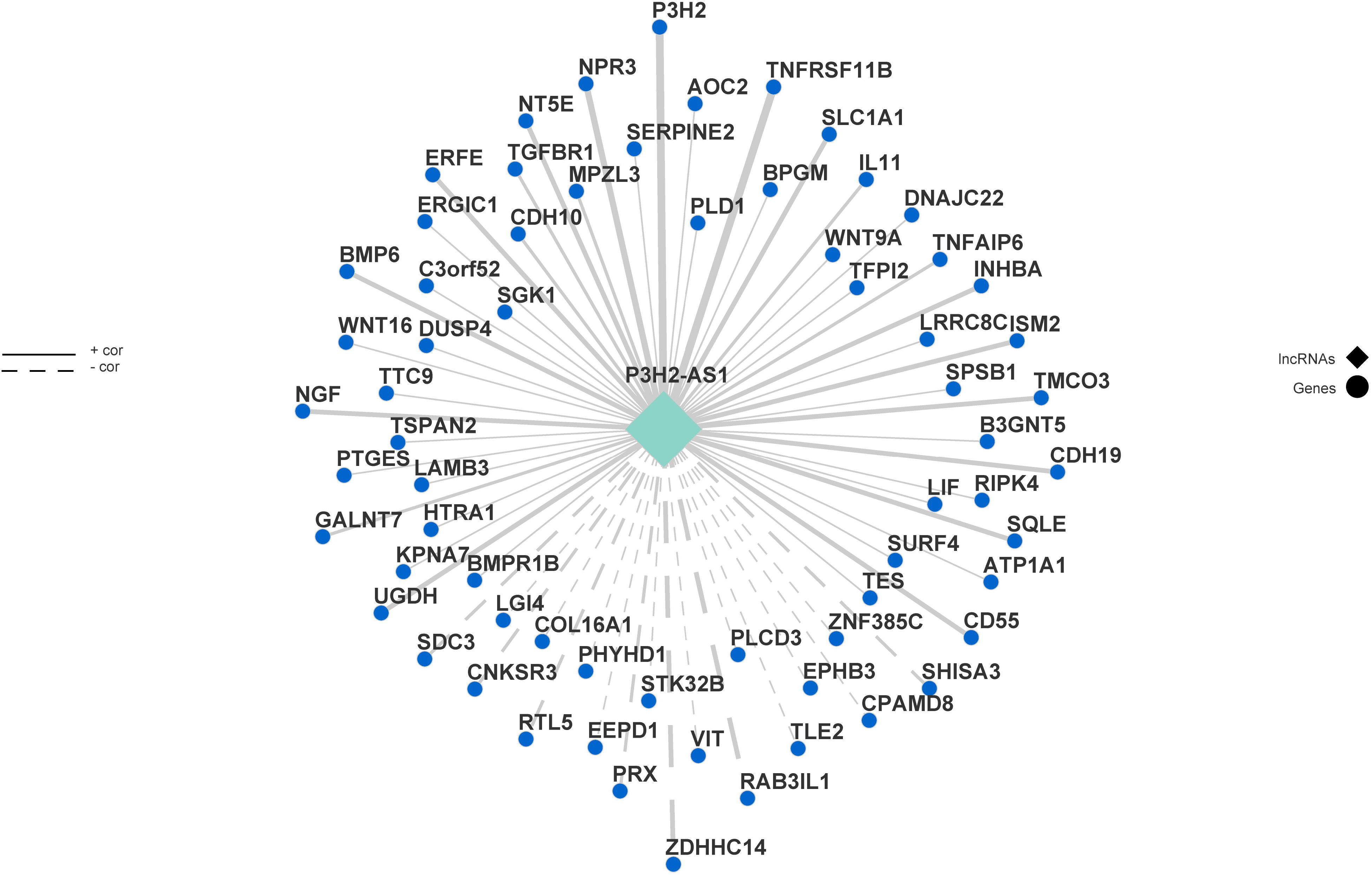
*P3H2-AS1*-mRNA co-expression network

## Notes

**Conflict of interests :** Not declared

## References

1. Goldring, S.R. and M.B. Goldring, Changes in the osteochondral unit during osteoarthritis: structure, function and cartilage-bone crosstalk. Nat Rev Rheumatol, 2016. 12(11): p. 632–644.

2. Tonge, D.P., M.J. Pearson, and S.W. Jones, The hallmarks of osteoarthritis and the potential to develop personalised disease-modifying pharmacological therapeutics. Osteoarthritis and Cartilage, 2014. 22(5): p. 609–621.

3. Ramos, Y.F., et al., Genes involved in the osteoarthritis process identified through genome wide expression analysis in articular cartilage; the RAAK study. PLoS One, 2014. 9(7): p. e103056.

4. Coutinho de Almeida, R., et al., RNA sequencing data integration reveals an miRNA interactome of osteoarthritis cartilage. Ann Rheum Dis, 2019. 78(2): p. 270–277.

5. Coutinho De Almeida, R., Y.F.M. Ramos, and I. Meulenbelt, Involvement of epigenetics in osteoarthritis. Best Practice & Research Clinical Rheumatology, 2017. 31(5): p. 634–648.

6. den Hollander, W., et al., Transcriptional associations of osteoarthritis-mediated loss of epigenetic control in articular cartilage. Arthritis Rheumatol, 2015. 67(8): p. 2108–16.

7. Jiang, S.D., et al., Long noncoding RNAs in osteoarthritis. Joint Bone Spine, 2017. 84(5): p. 553–556.

8. Pearson, M.J., et al., Long Intergenic Noncoding RNAs Mediate the Human Chondrocyte Inflammatory Response and Are Differentially Expressed in Osteoarthritis Cartilage. Arthritis Rheumatol, 2016. 68(4): p. 845–56.

9. Sun, H., et al., Emerging roles of long noncoding RNA in chondrogenesis, osteogenesis, and osteoarthritis. Am J Transl Res, 2019. 11(1): p. 16–30.

10. Liu, Q., et al., Long Noncoding RNA Related to Cartilage Injury Promotes Chondrocyte Extracellular Matrix Degradation in Osteoarthritis. Arthritis & Rheumatology, 2014. 66(4): p. 969–978.

11. Frankish, A., et al., GENCODE reference annotation for the human and mouse genomes. Nucleic Acids Research, 2019. 47(D1): p. D766–D773.

12. Jarroux, J., A. Morillon, and M. Pinskaya, History, Discovery, and Classification of lncRNAs. Adv Exp Med Biol, 2017. 1008: p. 1–46.

13. Gil, N. and I. Ulitsky, Regulation of gene expression by cis-acting long non-coding RNAs. Nat Rev Genet, 2020. 21(2): p. 102–117.

14. Villegas, V.E. and P.G. Zaphiropoulos, Neighboring gene regulation by antisense long non-coding RNAs. Int J Mol Sci, 2015. 16(2): p. 3251–66.

15. Uszczynska-Ratajczak, B., et al., Towards a complete map of the human long noncoding RNA transcriptome. Nature Reviews Genetics, 2018. 19(9): p. 535–548.

16. Ajekigbe, B., et al., Identification of long non-coding RNAs expressed in knee and hip osteoarthritic cartilage. Osteoarthritis Cartilage, 2019. 27(4): p. 694–702.

17. Dobin, A., et al., STAR: ultrafast universal RNA-seq aligner. Bioinformatics, 2013. 29(1): p. 15–21.

18. Pertea, M., et al., StringTie enables improved reconstruction of a transcriptome from RNA-seq reads. Nature Biotechnology, 2015. 33(3): p. 290–295.

19. Cunningham, F., et al., Ensembl 2019. Nucleic acids research, 2019. 47(D1): p. D745–D751.

20. Han, S., et al., LncFinder: an integrated platform for long non-coding RNA identification utilizing sequence intrinsic composition, structural information and physicochemical property. Briefings in Bioinformatics, 2018.

21. Love, M.I., W. Huber, and S. Anders, Moderated estimation of fold change and dispersion for RNA-seq data with DESeq2. Genome Biology, 2014. 15(12).

22. Ritchie, M.E., et al., limma powers differential expression analyses for RNA-sequencing and microarray studies. Nucleic Acids Research, 2015. 43(7): p. e47–e47.

23. Bomer, N., et al., Neo-cartilage engineered from primary chondrocytes is epigenetically similar to autologous cartilage, in contrast to using mesenchymal stem cells. Osteoarthritis Cartilage, 2016. 24(8): p. 1423–30.

24. Chen, K., et al., LncRNA MEG3 Inhibits the Degradation of the Extracellular Matrix of Chondrocytes in Osteoarthritis via Targeting miR-93/TGFBR2 Axis. Cartilage, 2019:p. 1947603519855759.

25. Zhang, Y., et al., LncRNA MALAT1 promotes osteoarthritis by modulating miR-150-5p/AKT3 axis. Cell & Bioscience, 2019. 9(1).

26. Tang, L.P., et al., LncRNA TUG1 promotes osteoarthritis-induced degradation of chondrocyte extracellular matrix via miR-195/MMP-13 axis. Eur Rev Med Pharmacol Sci, 2018. 22(24): p. 8574–8581.

27. Xing, D., et al., Identification of long noncoding RNA associated with osteoarthritis in humans. Orthop Surg, 2014. 6(4): p. 288–93.

28. Fu, M., et al., Expression profile of long noncoding RNAs in cartilage from knee osteoarthritis patients. Osteoarthritis and Cartilage, 2015. 23(3): p. 423–432.

29. Charles Richard, J.L. and P.J.A. Eichhorn, Platforms for Investigating LncRNA Functions. SLAS TECHNOLOGY: Translating Life Sciences Innovation, 2018. 23(6): p. 493–506.

